# Unexpected diversity of CRISPR unveils some evolutionary patterns of repeated sequences in *Mycobacterium tuberculosis*

**DOI:** 10.1101/2019.12.13.875765

**Authors:** Guislaine Refrégier, Christophe Sola, Christophe Guyeux

## Abstract

Diversity of the CRISPR locus of *Mycobacterium tuberculosis* complex has been studied since 1997 for molecular epidemiology purposes. By targeting solely the 43 spacers present in the two first sequenced genomes (H37Rv and BCG), it gave a biased idea of CRISPR diversity and ignored diversity in the neighbouring *cas*-genes.

We set up tailored pipelines to explore the diversity of CRISPR-cas locus in Short Reads. We analyzed data from a representative set of 198 clinical isolates as evidenced by well-characterized SNPs.

We found a relatively low diversity in terms of spacers: we recovered only the 68 spacers that had been described in 2000. We found no partial or global inversions in the sequences, letting always the Direct Variant Repeats (DVR) in the same order. In contrast, we found an unexpected diversity in the form of: SNPs in spacers and in Direct Repeats, duplications of various length, and insertions at various locations of the IS*6110* insertion sequence, as well as blocks of DVR deletions. The diversity was in part specific to lineages. When reconstructing evolutionary steps of the locus, we found no evidence for SNP reversal. DVR deletions were linked to recombination between IS*6110* insertions or between Direct Repeats.

This work definitively shows that CRISPR locus of *M. tuberculosis* did not evolve by classical CRISPR adaptation (incorporation of new spacers) since the last most recent common ancestor of virulent lineages. The evolutionary mechanisms that we discovered could be involved in bacterial adaptation but in a way that remains to be identified.

## Introduction

Since the rise of molecular biology, repeated sequences (CRISPR, IS, VNTRs) have been used to track relatedness between individuals (Garcia De Viedma and Perez-Lago, 2018). Indeed, they share two major features essential for diversity studies: ease of study, and rapid mutation rate (van Belkum, et al., 1998). In pathogens like *Mycobacterium tuberculosis* complex (MTC) they have been used for molecular epidemiology, complementing contact tracing, and/or identifying unsuspected links (Garcia De Viedma and Perez-Lago, 2018). In the last 5 years however, popularity of most repeated sequences has decreased first because they are larger than reads provided by Short Reads Sequencing, and second because of the generalization of Whole-Genome-Sequence availability and use of softwares analyzing Single Nucleotide Polymorphisms (SNPs) (Jajou, et al., 2018; Schurch, et al., 2010). In fact, some of these repeated sequences have sufficient variation to characterize them based on reads. The boom of Whole Genome Sequencing provides plenty of data to dig into for evolutionary studies and changes the way drug-susceptibility testing will be done in the future (Consortium, et al., 2018; Mulholland, et al., 2019). We will show in the case of CRISPR sequences how this diversity can reveal unexpected evolutionary patterns. We will show in addition that in the species of focus, namely MTC, there has been no new spacers acquisition for at least 5,000 years, *i.e.* no adaptative evolution in the common CRISPR terminology despite the presence of *Cas* genes.

CRISPR acronym stands for Clustered Regularly Interspaced Short Palindromic Repeats (Jansen, et al., 2002). They are characterized by repeats of 21 to 37 nt called Direct Repeats (DR) and the presence of unique sequences, called spacers, between each DR copy. Blocks of one DR and the following spacer has been termed Direct Variable Repeat (DVR) (Groenen, et al., 1993). CRISPR loci were first identified in *Escherichia coli* (Ishino, et al., 1987), their role in bacterial immunity was suspected in *Yersinia pestis* (Pourcel, et al., 2005), and later demonstrated in *Streptococcus thermophilus* (Barrangou, et al., 2007). Their presence has been detected in around 50% percent of eubacteria and 90% of archaebacteria (Couvin, et al., 2018; Grissa, et al., 2008; Grissa, et al., 2007; Grissa, et al., 2007). Various classes of CRISPR systems have been described (Makarova, et al., 2015). They all share the same mechanism of spacer acquisition, inserting part of a foreign sequence designated as *protospacer*, with a length similar to that of the repeats, next to the 5’ end of the locus. In *Salmonella enterica* for instance, the exploration of CRISPR diversity has shown that sequences including several DVR could be deleted, and that mutations could occur in spacers (Fabre, et al., 2012), however, the increased CRISPR dictionary as well as the restricted number of genomes sequenced reduced the possibility to have an extensive understanding of their evolutionary mechanisms.

*Mycobacterium tuberculosis* complex (MTC) is the agent of mammal tuberculosis, with human-adapted lineages being the most diverse and well spread. Its emergence and diversification dates back to at least 5,000 years old. There are six main and widely spread human-adapted sublineages referred to as L1 to L6 and an animal-adapted lineage (Coll, et al., 2014; Gagneux, 2012; Hershberg, et al., 2008), as well as a few rare and endemic human lineages (L7, L8, L0) (Blouin, et al., 2012; Ngabonziza, et al., 2019). Their diversity is being progressively unveiled through extensive WGS (Coll, et al., 2014; Palittapongarnpim, et al., 2018; Shitikov, et al., 2017).

*M. tuberculosis* reference clinical isolate H37Rv as well as most *M. tuberculosis* isolates carry a CRISPR locus together with a complete *cas* genes set of type III-A (Makarova, et al., 2015). Rare isolates lack part of CRISPR and or *cas* genes (Freidlin, et al., 2017). Partial analysis of the CRISPR diversity has been used since 1997 to explore the clinical isolates relatedness through a technique coined as « spoligotyping » (Kamerbeek, et al., 1997). In this technique, the presence of 43 spacers identified in H37Rv (n=35) or in *M. bovis* BCG (n=8) are looked for. This results in a barcode that can be easily shared and stored. Spoligotyping has led to the set-up of the first worldwide database for this pathogen counting today more than 111,000 patterns originating from 169 countries (Couvin, et al., 2018). The absence in some isolates of individual or consecutive spacers has revealed the possibility for small and large deletions of adjacent DVR (Brudey, et al., 2006; Filliol, et al., 2003). Large deletions proved good markers of tuberculosis diversification (Comas, et al., 2009; Kato-Maeda, et al., 2011).

Extensive MTC CRISPR structure has been previously explored in 19 *M. tuberculosis* clinical isolates belonging to EAI (L1), Beijing (L2), Euro-American (L4) lineages, 5 from animal species *M. bovis* and *M. microti*, and one *M. canettii* (van Embden, et al., 2000). This work showed that additional diversity exists in the form of DR variants, and duplication of DVR. It also documented the presence of insertion sequence IS*6110* in two different positions and orientations in L2 and L4 lineages. CRISPR diversity however remains unexplored in many sublineages as well as in major lineages such as L3, L5 and L6.

We recently set up a pipeline to reconstruct reliably CRISPR locus of *M. tuberculosis* (Guyeux, et al., 2019a). We selected Short Reads Archives (SRA) from the more than 60,000 available today to represent clinical isolates diversity and derived their CRISPR locus structure. The specific questions we tackled are: does MTC CRISPR locus contain additional spacers in addition to the 68 spacers ones described? What are the other patterns of diversity in CRISPR-Cas locus? What kind of underlying mechanisms of evolution can account for the observed diversity? Did the main lineages evolve similarly or are they CRISPR features specific of some lineages and/or sublineages? What is the most likely CRISPR sequence of tuberculosis most recent common ancestor (MRCA)?

## Methods

### Data collection

One hundred ninety-eight (n=198) Sequence Reads Archives obtained by paired-end sequencing with Illumina technology were selected from a local database of more than 3,500 genome sequences based on their representativeness of *M. tuberculosis* lineages (Guyeux, et al., 2019a). Namely, the following numbers of data were included for each lineage: 55 for Lineage 1, 20 for Lineage 2, 17 from Lineage 3, 60 from Lineage 4, 25 from Lineage 5, 7 from Lineage 6, 10 from Lineage 7, 1 from *M. bovis*, 1 from *M. caprae*, 1 from *M. microti*, 1 from *M. pinnipedii*. Data were downloaded as fasta files to decrease storage space as erroneous sequence will be ignored in the analytic steps.

### Identification and cataloging of CRISPR subsequences of interest

We first looked for spacer variants by searching for patterns made up of the last 12 nucleotides of most common DR sequence and later referred to as DR0 (Kamerbeek, et al., 1997), followed by 10 to 70 nucleotides, followed by the first 12 nucleotides of the DR0. The resulting subsequences were compared to the reference spacers to be declared either as a new spacer or a variant of a known spacer (for more details; see (Guyeux, et al., 2019a)). We then used this enhanced catalogue of spacers to find DR variants, in the same way as above. The new DRs thus obtained were used for a second phase of discovery of spacers, as described above.

To the collection of different spacers and DR, we added the following subsequences of interest to be discovered in the CRISPR loci:

1. the beginning and end sequences of IS*6110* and its reverse complement (40 bp each time);
2. those corresponding to *Rv2816c* (*Cas2* gene of the Cas locus) and *Rv2813c*, reputed to border the CRISPR locus;
3. the sequences found between these bordering genes and first or last DR;
4. the beginning and the end of each *Cas* gene;
5. sequences in the neighbouring genes (*Cas* or others) when these sequences were found besides an IS*6110* sequence during reconstruction –see below- (for more details; see (Guyeux, et al., 2019a)).

An extended version of these sequences of interest is presented in **Supplementary file 1**.

### Locus reconstruction

An automated contig building method based on De Bruijn approach was set up to reconstruct large fragments of the CRISPR. CRISPR with IS*6110* insertion could not directly be reconstructed as no read can overlap the full IS*6110* sequence (1,355 bp in length). Another reason for non-resolution of contigs is the existence of duplications: they lead to bifurcations in the de Bruijn graph. A specific search for duplications was included looking for patterns of the form sp(l)*DRX*sp(m), where l≥m (for more details; see (Guyeux, et al., 2019a)).

To facilitate the contigs concatenation, sequences were simplified by replacing each subsequence of interest by its name according to the catalogue described above. Final reconstruction taking into account IS*6110* insertions was performed manually. In some samples, contig reconstruction was confirmed by retrieving the identity of the spacer downstream the last spacer of a duplication. When one side of the CRISPR could not be automatically recovered for instance due to an IS*6110* insertion with a single end found in the catalog of CRISPR locus sequences, a stepwise manual search for the neighbouring sequences was performed until recovery of the other IS*6110* end. The 60nt sequence found nearby was labelled according to the gene it belongs to and its position, and it was added to the catalog of sequences of interest.

## Results

### 1. Exhaustive catalog of spacers in *M. tuberculosis* complex *stricto sensu*

We set up a method to identify not only variant of known spacers but also unknown spacers from *M. tuberculosis* CRISPR locus. Surprisingly, despite having explored more than 1,000 sequencing data (Guyeux, et al., 2019a), we found no new spacers as compared to the 68 described previously for *M. tuberculosis sensu stricto* (excluding *M. canettii* or the new L0 and L8 lineages) (van Embden, et al., 2000). The only new spacers that could be identified were found in *M. canettii* (data not shown). To identify whether this absence of new spacers could be due to a lack of sampling, we counted the cumulative number of spacers from the subset of isolates further described in this study upon 15 independent random samplings (**Figure 1**). We found that the 68 known spacers were all sampled after having examined from 3 to 25 isolates. Our sampling was therefore one order of magnitude above the one that seems necessary to be exhaustive.

**Figure 1.**
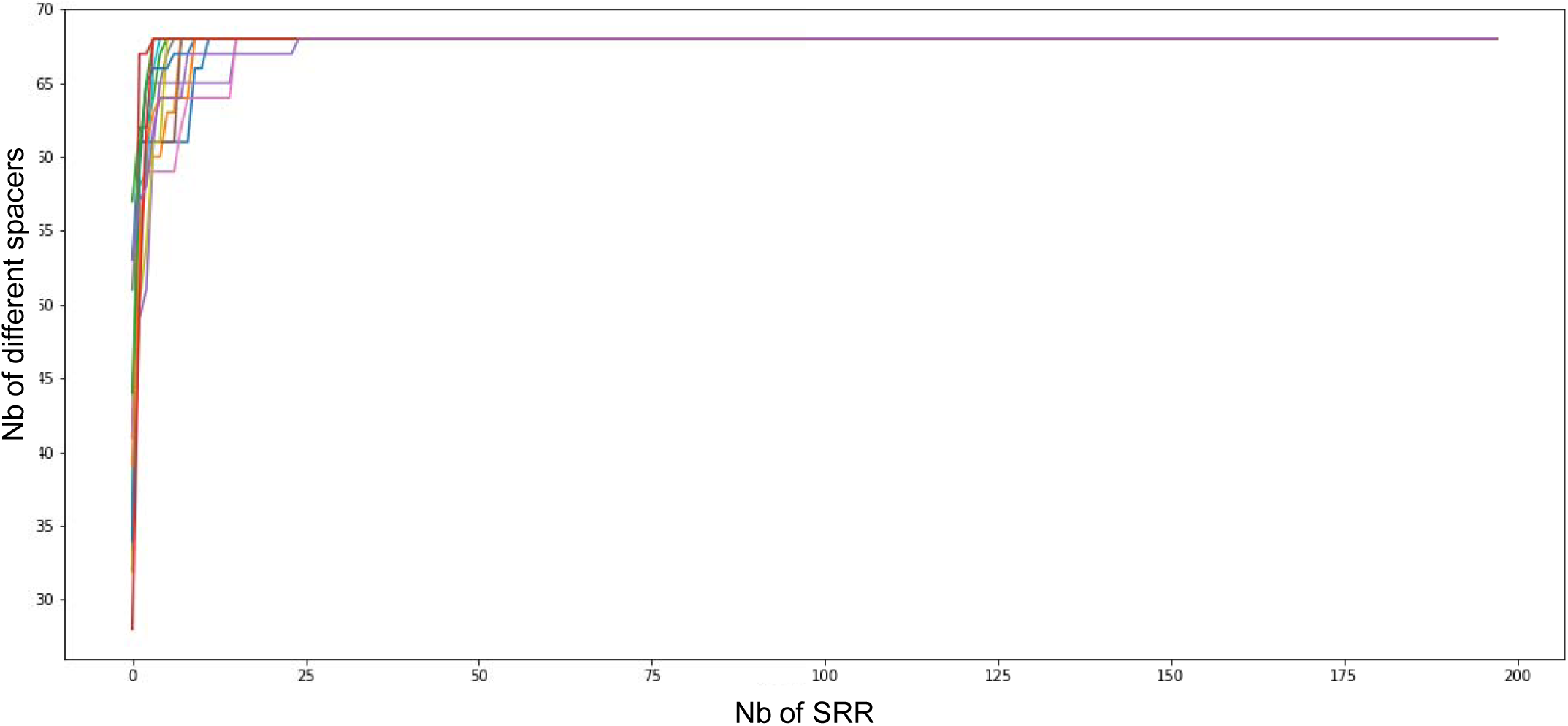
Cumulative number of spacers along random sampling of our database.

### 2. Global structure of *M. tuberculosis* CRISPR

We reconstructed the whole CRISPR loci for 198 clinical isolates representative of all *M. tuberculosis* diversity excluding *M. canettii*. CRISPR is almost always preceded by a complete set of *cas* genes, was followed by *Rv2813*, circumvented by one Direct Repeat sequence, DR0, at each of its border as can be seen for archetypal isolates from each Lineage (**Figure 2**). External DR0s are bordered by specific sequences, one of 48nt in length at the beginning of the locus, after *Cas2*, one of 148nt at the end of the locus, before Rv2813 (**Supplementary file 1**). These sequences are found in all isolates except in the case of large deletions (**Supplementary file 2**[IS*6110* sheet]). Most of the times, the CRISPR-Cas locus includes one IS*6110* copy as in the first isolate presented in **Figure 2** belonging to L1.1.1.6 (ERR751749), but it can go up to three copies or down to zero (**Supplementary file 2**[IS6110 sheet]). No other type of insertion sequence was ever discovered inside the region (data not shown).

**Figure 2.**
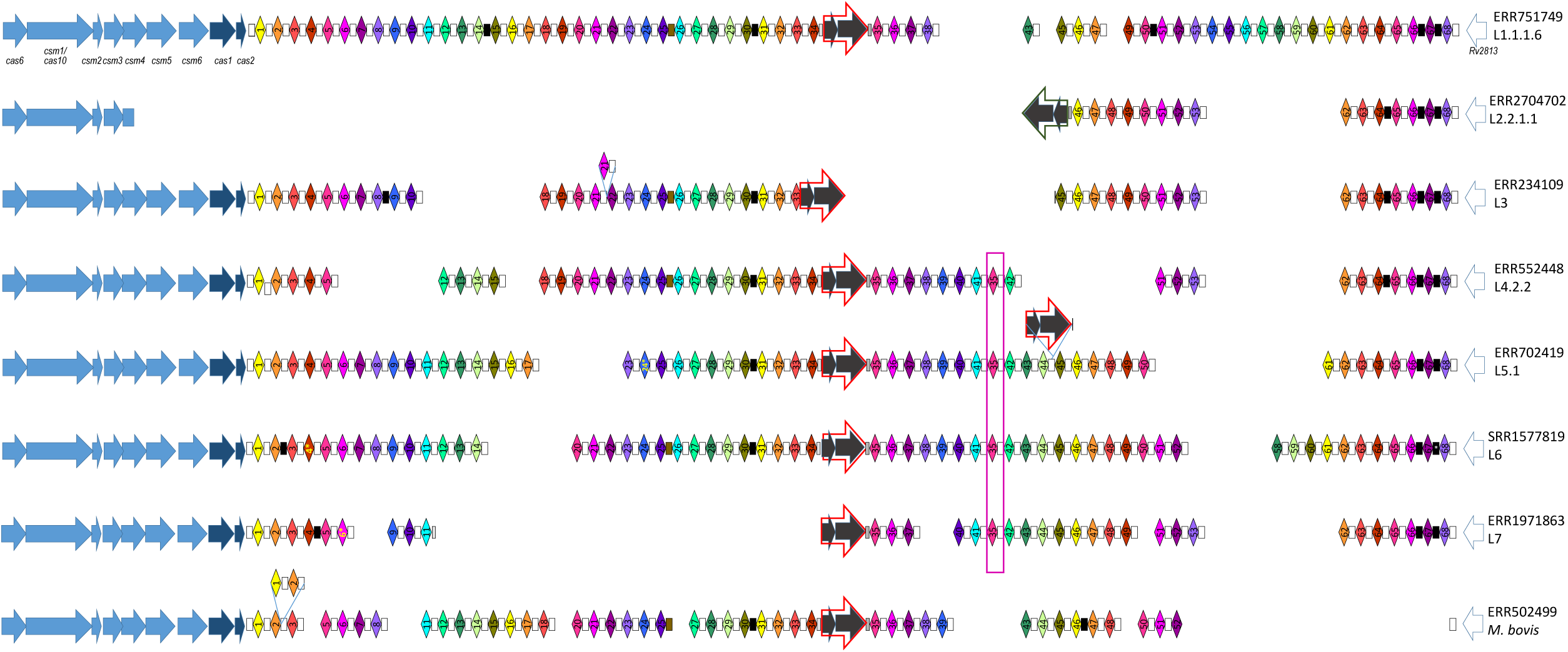
CRISPR-Cas locus reconstitution for one archetypal isolate of each lineage. *Notes common to fig. 2 and 3:* Arrows indicate genes. Diamonds indicate spacers. Boxes indicated Direct Repeats (DR). Width of spacers are DR has been articially expanded for clarity. The pink empty box highlights a duplicated spacer at an unexpected position (not in tandem). Color codes for genes (arrows): light blue: *cas* genes involved in immunity (interference); dark blue: *cas* genes involved in adaptation; green : IS*6110* genes (transposase and hypothetical protein); white: other neighbouring gene of unknown function. The color of spacers was attributed randomly to facilitate visual exploration but spacers of the same color have no link except if they carry the same number. Direction of CRISPR-Cas locus is antisense as compared to H37Rv genome orientation, so that all *cas* genes are annotated with a c: *cas6* is Rv2824c and *cas2* is Rv2816c. Genes forming the IS*6110* sequence are sometimes in the sense and sometimes in the antisense direction. Between spacers 34 and 35 as in H37Rv, there are in the antisense direction and therefore are referred to as Rv2815c and Rv2814c. Several DRs are truncated. Between spacers 34 and 35, IS*6110*c is preceded by a sequence close to rDra, corresponding to the 19 first nt of DR0 (shown in light grey), and is followed by a sequence close to Drb (referred to as DRb1) corresponding to the 20 last nt of DR0 (darker grey). These two sequences therefore share the CCC sequence in the middle of DR0. They are also found around the IS*6110*c sequence of L7 isolates. A similar case is true in L5.1 ERR7022419 clinical isolates. Around the IS*6110*c copy in ERR234109 (L3), the preceding spacer is slightly truncated (sp33, only its first 27 nt), and there are only the last 4 nucleotide of the DR0 before the nest spacer (sp45). When a DR0 borders a deletion, we chose to represent it in most of the cases at the beginning of the deletion, although choosing the end of the deletion would have been equally relevant. Mutated DR are indicated in black. They are not the same from one position to the other, but variants at the same location are the same except for the DR between spacers 67 and 68 that harbors a second variant solely in L6 and is therefore indicated by a star (see Supplementary file 3).

The spacer sequences as well as those of the DR are always found in the same direction. Their order of succession is usually the expected one (the order of natural integers) although, as described below, various particular situations arise, for instance in case of duplications (**Supplementary file 3**). Duplications are identified not only by the order of successive spacers, but also by the relatively higher number of reads corresponding to the duplicated spacers. For instance, in an isolate belonging to L1.1.1.8 (ERR718201), while most spacers were found on an average of 27 reads, spacers 14 to 21 are found in 56 reads on average, which is approximately twice as much (**Figure 3**). A notable exception in this isolate is spacer 16 that is found in only 31 reads. This however matches the fact that spacer 15 is half of the time followed by spacer 16 and the other half by spacer 17: in one of the two spacer 14-spacer 21 region, DVR16 has been deleted (**Figure 3**).

**Figure 3.**
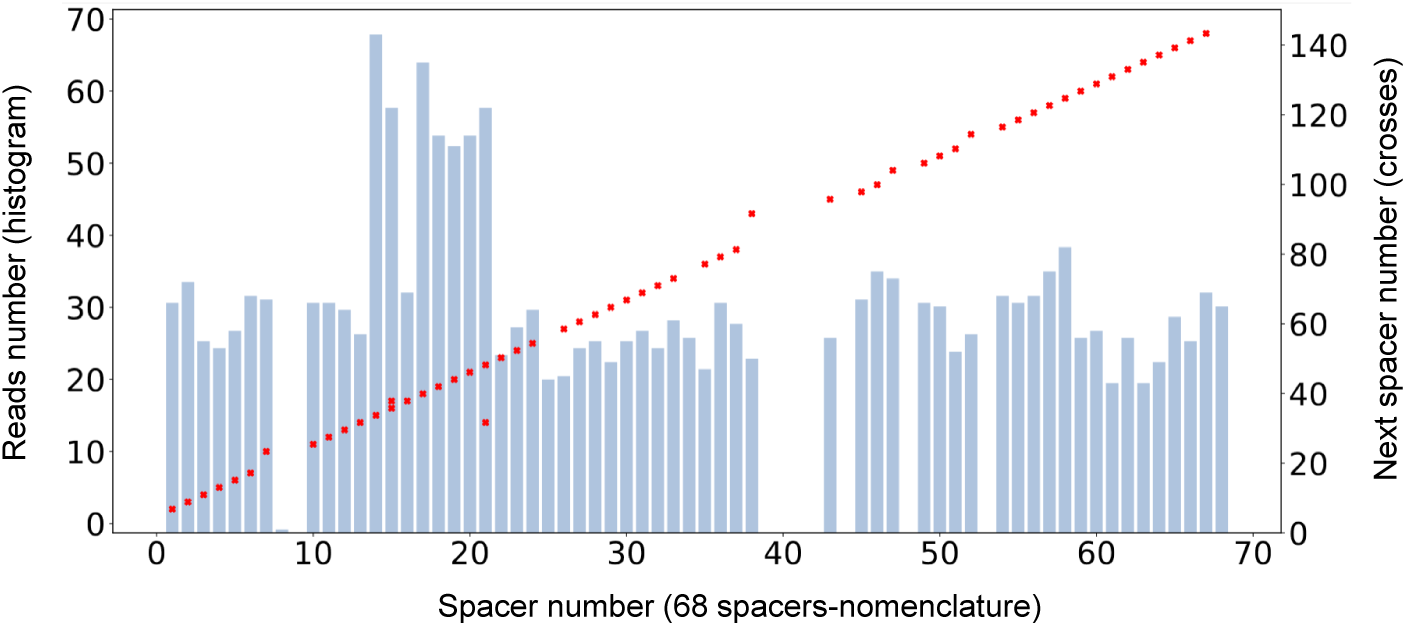
Proof for spacers 14-20 duplication in isolate ERR718201. Reads number as a function of spacer number is shown in blue. The number of the following spacer is shown in red (crosses).

Duplications occur in tandem most of the time. For instance, a second DVR21 is found after its normal copy in L2 isolates such as ERR234109, and an additional tandem DVR1-DVR2 is found downstream the standard pair in *M. bovis* ERR5022499 (**Figure 2**). Other examples include DVR32 in ERR234197 (L1.1.3.1), DVR39 in ERR234248 (L2.1). This can be seen directly in the Illumina sequences, for instance for ERR234248, where many reads contain the end of 39, followed by a DR0, followed by the beginning of another 39, which has no chance of happening, in such a repeated way, by chance due to random reading noise. A notable exception to the natural order of succession of spacer is the case of the spacer 35, which can be found in the following two places: between 34 and 36 on the one hand, and after 41 on the other hand (**Figure 2, Supplementary file 4**). Consequently, in most cases, although this is not the case of H37Rv and related isolates, there are two copies of 35.

Another important and widely representative characteristic of MTC CRISPR locus is the presence of the IS*6110* copy referenced in (Kamerbeek, et al., 1997) and that shares the same orientation than the CRISPR, *i.e.* corresponding to a IS*6110*c (**Figure 2**).

### 3. Punctual variants in *M. tuberculosis* CRISPR

Regarding intra-spacer diversity, we identified 20 spacers that harbored at least two variants, and concerned 48 (24%) out of the 198 isolates explored (**Supplementary file 2**[spacer sheet]). These variants consisted mainly of SNP, although a deletion was found in spacer 24 in another dataset (genome ERR702419, lineage 5, data not shown). Interestingly, some of these variants are characteristic of specific lineages. For instance, a variant of spacer 38 is found in all isolates of lineage L1.1.1, one mutation is found in spacer 4 in all L6 isolates to which an additional one sometimes adds resulting in two possible variants. Two variants of spacer 6 characterize the endemic Abyssinian L7 isolates (**Figure 2, Supplementary file 5**). The frequency of spacer variants in L2-L3-L4-L7 was relatively low (6 independent variants detected in 107 isolates, ∼5%), as compared to L1 lineage (11 independent variants out of a selection of 55 isolates, ∼20%) and lineage gathering animal isolates and L5 and L6 (7 independent variants for 34 isolates, ∼20%).

Between two spacers, we have most of the time the DR0 sequence referenced in (van Embden, et al., 2000). However, this rule is incomplete and not general. Punctual variants were identified. First of all, between spacers 30 and 31, there is always, whatever the lineage, a sequence that we coined DR2 and that has one punctual mutation as compared to DR0 (see sequence in **Supplemental file 1**). Similarly, there is always a DR4 variant repeat between spacers 66 and 67, and again a DR5 variant between spacers 67 and 68. This is true for all lineages, with the notable exception of a sublineage of L6, which has yet the DR10 variant (**Figure 2, Supplementary file 2**[DR sheet]). Then, other types of variations were identified. For instance, between spacers 25 and 26, there are always only the last 24 nt of DR0 (a sequence we name DRb2). Around the central IS*6110*c, between spacers 34 and 35, the DR0 is split into two subsequences rDRa1 (upstream) and DRb1 (downstream). As expected due to IS*6110* insertion characteristics, the concatenation of these two sequences is 3nt larger than DR0 since 3 additional cytosines are present at each end of the insertion (Gonzalo-Asensio, et al., 2018; Thierry, et al., 1990). Yet, in a L5.1 isolate (ERR702419) where IS*6110*c inserted downstream spacer 44, IS*6110*c is preceded by the first 35 nt of DR0 and followed by its 6 last nt, so that the duplicated target was this time 5nt in length (data not shown).

Some variants are shared over several but not all lineages or sublineages. For instance, DR6 is found between spacers 64 and 65, in all genomes of lineages L2 to L4, and only in those; DR10 is found between spacers 67 and 68 in L6. Similarly, the DR1 variant is found between 14 and 15 only in Sublineage L1.1.1, and never in Sublineage L1.1.2, or in any other lineage. These findings are consistent with *M. tuberculosis* phylogeny and allow to infer that the mutation in L1.1.1 occurred shortly after separation from the rest of the other L1 sublineages.

Other punctual variants affect a single isolate (**Supplementary file 2**[spacer and DR sheets] for isolates affected, **Supplementary file 1** for their sequence). Each time, the size of the DR is respected (no indel, only the single nucleotide polymorphism) except for one case where a longer DR was found (data not shown). Altogether, these variants occurred all over the locus with no clear preferential subregion (**Supplementary file 6**).

### 4. Large scale variations and IS*6110* copies

Large scale variations included on one hand deletions and, on the other hand, duplications. It should be noted that, at this stage, no inversion has been detected in MTC CRISPR.

Large-scale deletions were observed throughout the lineages, such as the one characterizing L2.2/Beijing sublineage that covers parts of *csm4* to an IS*6110* just before spacer 46 (#36 in the old nomenclature). As in the case of this specific deletion, many deletions were flanked by an IS*6110* insertion: the deletion between spacer 33 and spacer 45 in L3 isolates ERR234109, and the deletion between spacer 11 and spacer 35 in L7 isolates ERR1971863 (**Figure 2**). To infer potential intermediates for these deletions, we searched for clinical isolates related to the one carrying deletions, and harbouring several IS*6110* sequences. We found such evidence in Sublineage L4.1.2.1 (Haarlem sublineage). In this sublineage, a first set of isolates carry a 7 DVR-deletion adjacent to an IS*6110* copy, namely between spacers 34 and the second copy of spacer 35 (for instance in ERR234259). A second set of clinical isolates (SRR5073877 and ERR552680) harbours two IS*6110* copies, respectively the well-known one in the DR between spacers 34 and spacer 35, and another one in the DR between spacer 41 and the second spacer 35 (**Figure 4**). Interestingly, the borders of IS*6110* insertion in ERR234259 corresponded well to the external borders of the two IS present in SRR5073877 and ERR552680. The left border consisted in the 17 first nt of DR0 (2nt less only than the rDRa1 in the classical position), and the right border was the exact same 33-last nucleotides of DR0 than the one found at the right of the second insertion in SRR5073877 and ERR552680. The CRISPR version with the two copies shares many features with that carrying the deletion, suggesting that it could correspond to its ancestral stage of evolution (**Figure 4**). The similar observation in L4.1.2.1 was made independently in a study performed in Hanoi (Maeda, et al., 2020).

**Figure 4.**
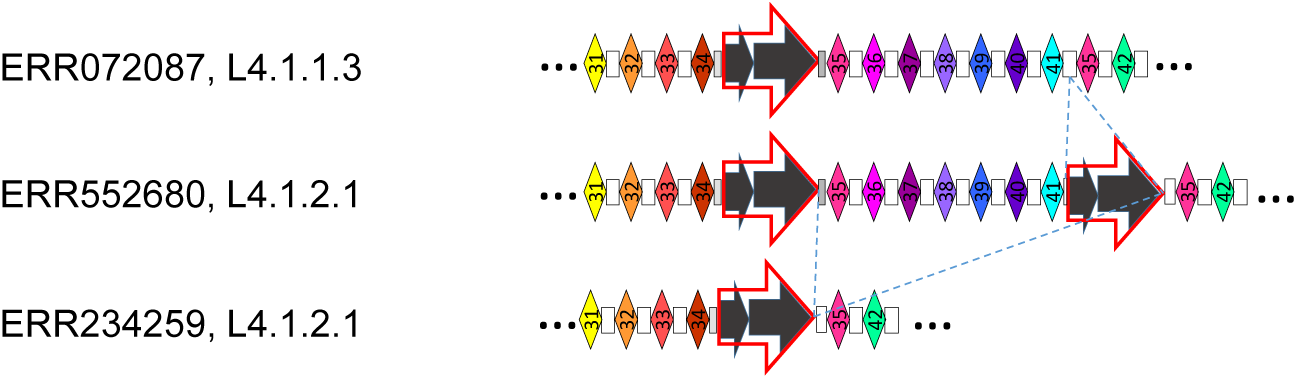
CRISPR substructures of related isolates illustrating deletion by recombination between IS*6110* copies. ERR072087 with one single copy with all spacers in the subregion of interest likely harbors the most ancestral structure. ERR552680 with two copies and all spacers likely represents an intermediate state after a new IS*6110* insertion. ERR234259 with a single copy and loss of spacers likely emerged due to the recombination between the two copies present in ERR552680.

**Figure 5.**
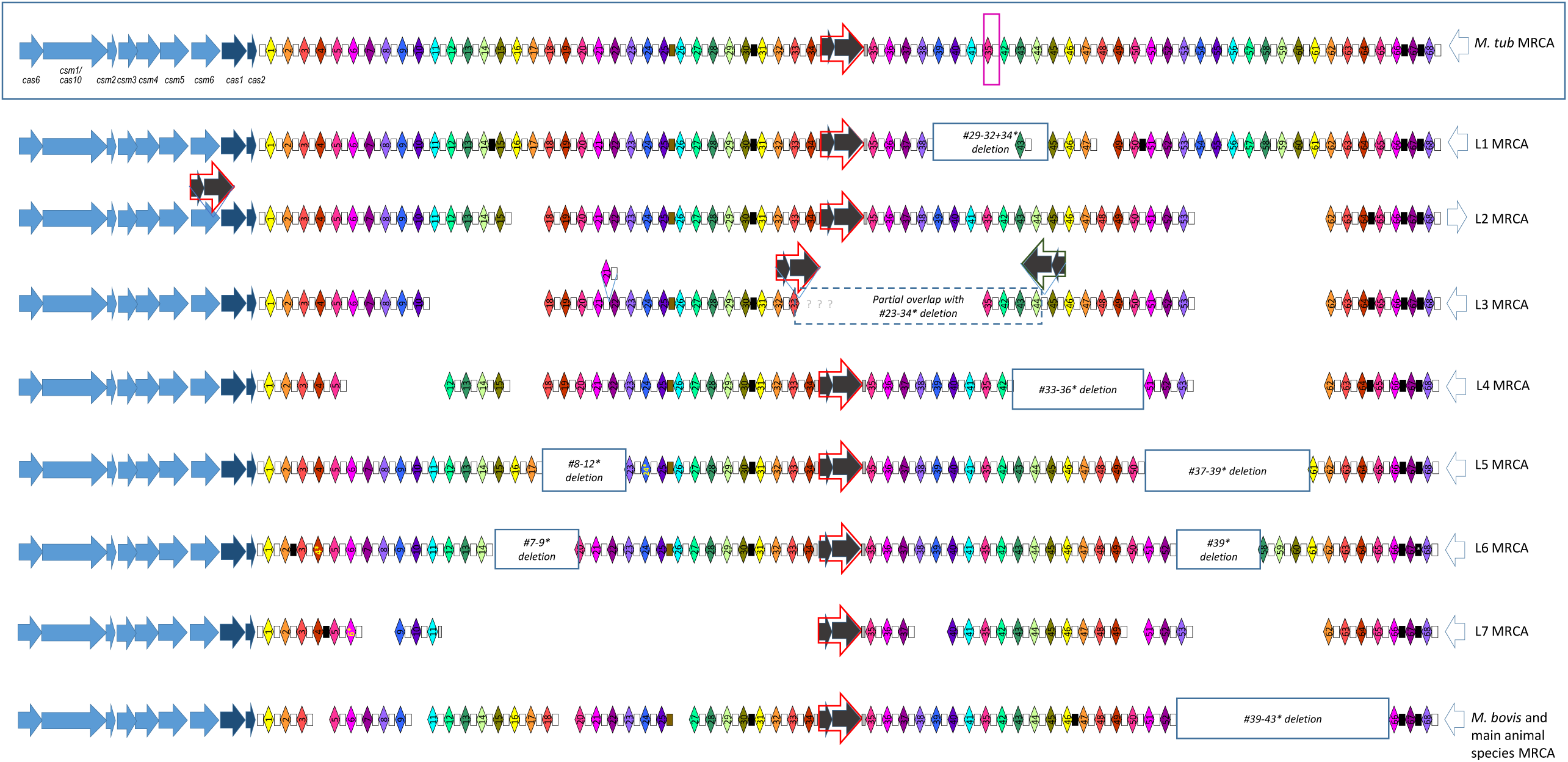
CRISPR-Cas locus likely structure of each lineage MRCA. The proposed structure was designed by a parsimonious approach based on the CRISPR structure of the 198 clinical isolates fully characterized in Supplementary file 3 (See also notes common with Fig. 2).

These large scale deletions involved *cas* flanking genes in 23/198 (12%) of isolates, with two different borders in L2 isolates, two others in L4 and a third one in L3. In contrast, a single case was observed that affected *Rv2813* (**Supplementary file 2**[IS6110 sheet]). We further explored this asymmetry using SITVIT2 2019 database (n=3852 SITs): 290 SITs harbored a deletion of spacer #1 (DVR2 in the new nomenclature) against 117 SITs with a deletion of spacer #43 (DVR65 in the new nomenclature), *i. e.* three times more deletions on the *cas* genes side.

### 5. Likely MRCA CRISPR of *M. tuberculosis*

All variations we observed were concordant with the phylogeny of *M. tuberculosis*. We could thus infer the most likely structure of CRISPR locus of *M. tuberculosis* complex *sensu stricto* (without *M. canettii*), as well as its structure in all MRCA lineages. We found that global MRCA likely carried a full set of *cas* genes, a CRISPR with 69 spacers (the 68 spacers of different sequences + the repetition of spacer 35) interspersed mostly by DR0 except between spacers 25 and 26 (DRb2), spacers 30 and 31 (DR2), spacers 66 and 67 (DR4) and spacers 67 and 68 (DR5). An ancestral and central IS*6110*c was inferred to lie at the same place as the one occupied in H37Rv, i.e. between spacers 34 and 35 (**Figure 4**). A deletion of DVR 54 to 61 characterized MRCA of lineages 2, 3, 4 and 7, which is not documented in the classical form of the spoligotype as these spacers are not belonging to its set of 43 spacers. Other deletions corresponded to the ones found in spoligotype-43 format and used to define main sublineages. For instance, the deletion of spacers #33-36 in the old nomenclature for L4/Euro-American lineage (previously referred to as T family) corresponds to the deletion of DVR43 to 50. Another example is the deletion of spacers #29-32, presence of spacer #33 and absence of spacer #34 characteristic of Lineage 1 (previously referred to as EAI) (Filliol, et al., 2003) that corresponds to the deletion of DVR39 to 42, presence of DVR43 and absence of DVR44 (**Figure 4**). Only L2 MRCA did not carry the well-known signature of Beijing isolates as L2 includes not only the Beijing L2.2 sublineage but also the L2.1 proto-Beijing sublineage (Shitikov, et al., 2017). Interestingly, this ancestor harbors an IS*6110* insertion in one *cas* gene (namely *csm6*) but not at the border of the classical Beijing deletion. It also lacks DVR16 and DVR17.

## Discussion

Thanks to our new Sequence Reads Archive-based genomic analysis pipeline, we explored the *M. tuberculosis* CRISPR sequences diversity in 198 clinical isolates representative of the MTC excluding *M. canettii*, which deserve new specific studies (Supply, et al., 2013; van Soolingen, et al., 1997). These data show that *M. tuberculosis* CRISPR locus can contains at most 69 spacers (68 + one duplication), is not prone to inversions, evolves by duplication and deletions through recombination between DR, but also and primarily through insertion/deletions implicating IS*6110*, by homologous recombination, and independently of lineage. We detail below the support for these different kinds of mutations and inferences that can be drown concerning the functionality of CRISPR-Cas locus.

### Evolutionary mechanisms of MTC CRISPR locus expansion

Despite the absence of acquisition of new spacers, MTC CRISPR locus is of relative long size in many isolates (for instance, 4,589 nt between Rv2813 and Rv2816c/*cas2* in H37Rv). This relates to its ability tocontinue to expand using mechanisms other than classical CRISPR adaptation.

A first mechanism of MTC CRISPR size expansion, when considered as the distance between its two borders, is the integration of IS*6110* insertion sequences (1,355 bp). The most frequent insertion is found between spacers 34 and 35 as in H37Rv genome. Other IS*6110* insertions were found along the whole MTC CRISPR locus, with up to two insertions in the CRISPR locus and three when considering the whole CRISPR-Cas locus. Other similar IS Sequences right next to or farther away, might be responsible for other homologous recombination mechanisms involving CRISPR.

The second CRISPR expansion mechanism identified in this overall review concerns duplications of DVR (DR + spacer). These duplications are of two main types. First of all, duplications can concern single DVR and place in tandem which was observed in 11 independent cases throughout our 198 samples. This type of tandem duplication concerns also several adjacent DVRs such as DVR1-2 in *M. bovis* or DVR14-15-16-17-18-19-20 in L1.1.1.7. Such multiple DVR duplications were observed 5 times in our sample, so that in total 16 independent events of tandem duplications were observed. The second type of duplications concerns DVR that are far away from their original position, a type we call “rearrangement duplications”. This first concerns DVR35 located between DVR41 and DVR42 as already mentioned above and supposedly in MTC MRCA CRISPR. Other examples include a second copy of DVR3 found between DVR12 and 13 found in ERR036187 (L4.3.4.1), while in ERR234197 (L1.1.3.1), there is an additional copy of DVR38 between DVR55 and 56. In one instance, this concerned several adjacent DVRs: a second copy of DVR50-51-52-53 is found between DVR3 and 4 in ERR2245409 (L3.1.1). Altogether, this made a total of 4 independent rearrangement-duplications. The fact that rearrangement duplications are less common than standard duplications suggests that they occur less frequently and/or that they are less stable. If the stability of rearrangement duplications was low, there should be several cases of deletions between the two copies of DVR35 as they were likely already present in MTC MRCA. Yet, we observed no case where a deletion concerned solely the DVR between these two copies.

Overall, the proportion of genomes containing either several copies of IS*6110* or a duplication of one of the forms listed above is important, showing that MTC CRISPR is much more variable than what could be derived from a standard 43 spacers spoligotyping analysis. This is true not only for the *in vitro* but also for the *in Silico*-based acquisition of the spoligotype, as the blast procedure used in the current analytic tools (Spolpred, SpoTyping) only provides information on the presence or absence of a given spacer: there is nothing quantitative or location-related in these approaches (Coll, et al., 2012; Xia, et al., 2016). Hence, on one hand, the representation of the CRISPR locus through a simple barcode of presence/absence of individual spacers hide these quantitative and localization information, wheras on another hand, a more extensive description of the CRISPR locus including duplications, insertions, point mutations, provides useful information to classify and/or cluster clinical isolates. Such an information is advantageously correlated with the current SNPs based taxonomical system of MTC genomes and enhance our understanding of isolates evolution (Coll, et al., 2014; Palittapongarnpim, et al., 2018; Shitikov, et al., 2017; Stucki, et al., 2016).

### Combined Mechanism of CRISPR locus reduction: how does IS*6110* contributes to the evolution of CRISPR locus in MTC?

In addition to the undeniable expansion mechanisms mentioned above, CRISPR reduction mechanisms also coexist, which -to some extent-explain some of the spacer block deletions in MTC spoligotypes.

The first potential mechanism is the simple loss of spacer, for instance by recombination between two adjacent DRs. For instance, clinical isolate ERR1203071 of L4.8 lacks spacer 1. In place, it harbors a one nucleotide variant of the beginning sequence, a DR0 and spacer 2. The principle of parsimony here tends to suggest that a recombination between the DR0 bordering spacer 1 led to this genotype. The same kind of recombination seems to occur on slightly higher number of DVR such as the DVR54-DVR61 deletion typical of L2-3-4-7. Recombination between perfect DR would be favored compared to mutated DR.

We can know confidently argue that the second highly frequent mechanism, that is at play for the largest suppressions of consecutive spacers, is an IS-linked three steps mechanism: (1) insertion or prior presence of a first copy of IS*6110* (for instance that after spacer 34), (2) insertion of a second IS*6110* copy at another location (e.g. in *csm6* in the ancestor of L2, also seen in SRR1710060, see **Supplementary file 2**), and (3) recombination between the two IS*6110* copies. This IS-mediated mechanism, that has been described in previous studies is a general mechanism, i.e. it happens independently of lineage and is the responsible of IS*6110* convergence of IS copy numbers (Roychowdhury, et al., 2015). The final result is the change from *x* to *x-1* copies of IS*6110*, with the loss of all spacers between the two copies. This mechanism can be observed independently of lineages, for example, in lineage 4, in Haarlem (4.1.2.1): L4 ancestor has a single copy of IS between 34 and 35, then a second copy occurred in the ancestor of Haarlem L4.1.2 isolates as seen in ERR552680, between 41 and 35, and finally a deletion occurred leading to the loss of spacers 35 to 41 for some isolates such as ERR234259. It therefore seems reasonable to think that after the insertion after spacer 41, this copy of IS*6110* has recombined with the one upstream of spacer 35. This mechanism is also at work elsewhere in the Haarlem isolates between *csm5* and spacer 34 and between *csm5* and spacer 41 (**Supplementary file 2**).

IS*6110* insertions can take place in spacers or in DR and it is not necessary for an IS to be in a DR to be able to recombine. For instance, in many L4.3 (LAM) clinical isolates where spacers 31 to 34 (#21-#24) are missing, the successive sequences of interest are: the beginning of spacer 31 (#21), an IS*6110*c, DRb1 and spacer 35. The last three sequences of interest are found in the exact same order in undeleted isolates such as H37rv. This suggests that an IS*6110* copy was first inserted at the end of spacer 31, and that it later recombined with the one located between spacers 34 and 35. This recombination did not modify the flanking sequences.

The orientation of the two IS*6110* copies that recombined cannot always be derived due to the lack of the ancestral versions. Still in several cases, we could identify isolates related to the deleted ones, that carry the two IS*6110* flanking the future deletion. This is true for the IS*6110* insertions having led to the deletion described in **Figure 4**. In that case, both insertions were in the reverse sense as compared to H37Rv orientation and can be called IS*6110*c. In another case, the isolate with two IS*6110* insertions is SRR5073887 (L4.4.1): it carries not only the standard IS*6110*c insertion between spacers 34 and 35 but also an IS*6110* insertion in the sense direction at the 439^th^ nt of *csm6*. The deletion in ERR2653229 (also L4.4.1) flanked by the beginning of *csm6* and DRb1 and spacer 35 with a sense IS*6110* sequence in its middle (**Supplementary file 2** [IS6110 sheet]) likely occurred through the recombination of these two IS although they lie in opposite orientations. This phenomenon was recently observed in several cases of IS*6110* mediated deletions in L2 (Shitikov, et al., 2019).

### Variants and problems in spoligotyping

How does the sequence diversity impact spoligotyping data? When performed *in vitro*, spoligotyping consists first in the amplification of the CRISPR locus using primers facing the outside of DR region, referred to as DRa and DRb, and second in the hybridization to probes attached at a specific position on a membrane or another support. CRISPR sequences variants may reduce the efficiency of the process, whether at the amplification or at the hybridization step. The presence of intermediate signals in spoligotyping or discrepant results between *in Silico* and *in Vitro*-based spoligotypes has been documented by several authors (Abadia, et al., 2011; Meehan, et al., 2018). We looked for intermediate signals corresponding to variants. In the case of L6 clinical isolates that carry a variant of spacer 4 (spacer 3 in spoligo-43 nomenclature), we found no evidence of such report in the literature and in our own data (data not shown). The same was true for spacer 38 (spacer 28 in spoligo-43 nomenclature) found in L1.1.1 clinical isolates even if the mutation is relatively central in the probe (**Supplementary file 5**).

### Asymmetric variations affecting of MTC CRISPR-Cas locus

As described above, we identified punctual nucleotide mutations, duplications, IS insertions and deletions along CRISPR-Cas locus. CRISPR are oriented loci that acquire new spacers at the 5’ end relative to their transcription direction (Barrangou, et al., 2007; Makarova, et al., 2018). It may therefore be expected that variations do not affect symmetrically this locus. To explore and understand the consequences of this possibility, it is important to identify the orientation of the CRISPR locus in question. Using RNAseq data on H37Rv, Wei et al. showed that transcription occurs from spacer 1 towards spacer 68 (Wei, et al., 2019). We independently confirmed this observation by the exploration of independent RNAseq data from (Ignatov, et al., 2015; Rodriguez, et al., 2014) (Refregier et al. unpublished results). The orientation presented in this study is thus the functional one. According to classical CRISPR expansion mechanism, the introduction of new spacers occurs at the 5’ end of the locus, so that the most ancient DVR lies at its 3’ end.

In contradiction with the remarkable feature that most ancient DR carry mutations in all isolates, no subregion exhibited a significantly higher punctual mutation rate (**Supplementary file 6**). The fact that the most ancient part of CRISPR locus does not carry a significantly higher number of punctual mutations as compared to parts that are more recent (spacer block deletions), may suggest that the time during which the locus expanded from spacer 68 to spacer 1 may be negligible as compared to the time between MTC MRCA and present, or that the CRISPR locus was transferred by lateral gene transfer in one single block from another environmental organism. Alternatively, the time of CRISPR locus expansion could have been quite long, however the pace of CRISPR locus SNPs mutations acquisition was very slow because of an extremely slow pace of MTC transmission. Demography and genetic drift could have been much more important for MTC evolution than selection in human populations (Pepperell, et al., 2010). Yet, the presence of mutations in several DR at the 3’ end of the locus could also play a role in its stability.

In contrast, we detected an asymmetry concerning the loss of flanking sequences: it was apparently more frequent to have a loss of the beginning sequences of CRISPR, on the side of the *cas* genes (several independent isolates from L2 and from L4) than to have a loss of the ending sequences, i.e. on the side of *Rv2813*. All deletions implicating flanking sequences were bordered by an IS*6110* sequence. Altogether, the asymmetry in deletion suggests either a more crucial role of the end of the CRISPR *i.e.* of gene *Rv2813* and/or its neighbors, or asymmetric mechanisms favoring deletion on the *cas* gene side. This second possibility relates to IS*6110* insertion frequency as IS are always involved in large deletions. Saying that IS*6110* insertions are more likely on the *cas* gene side suggests either their lower impact on bacterial fitness, or a DNA superstructure that would favor IS insertions. Other IS exist in the genome that could also insert in a favorable region. Their presence in CRISPR region would be a sign that it is an integration hot spot. However, our script was designed to look only for insertion in *cas* gene that also lead to a deletion in the CRISPR in at least one of the explored sample. IS other than IS*6110* cannot lead to any deletion. Nevertheless, even if our script may have overlooked non-IS*6110* insertions, we did not encounter it in around 500 randomly sampled genomes. The question of *cas* gene locus being an integration hotspot of IS sequences needs other studies to be completely solved.

### Functionality of MTC CRISPR-Cas locus

CRISPR-Cas loci are involved in two mechanisms: 1) adaptation by the integration of new spacers, usually taken from foreign DNA, at the 5’ end of CRISPR with the help of Cas1 and Cas2 proteins, and 2) immunity by the transcription of CRISPR locus, processing with the help of Cas6 protein in the case of type III-A CRISPRs, and degradation of DNA and/or RNA carrying *protospacers*, with the help of the crRNP (CRISPR RiboNucleoProtein complex), a complex involving the crRNA and other Cas proteins. By exploring the diversity of many genomes at the CRISPR locus, we are able to infer the effectivity of adaptation processes. Regarding immunity, we can only state whether the necessary genes are present or not.

In the whole *M. tuberculosis* complex *sensu stricto*, we could find only the 68 spacers already present in the MRCA (van Embden, et al., 2000). We found no evidence that a single clinical isolate has acquired a new spacer in the course of MTC evolution. This seems particularly surprising as most currently spreading isolates apart those from L2 still carry the full set of *Cas* genes including *Cas1* and *Cas 2* involved in CRISPR adaptation in other type III-A systems. This could be due to a mutation in *M. tuberculosis* ancestor that has abolished *Cas1* and/or *Cas2* functionality in the ancestor. Another reason could be that MTC, given its intra-cellular life-style, does simply not have the chance anymore to encounter foreign DNA such as phages or plasmids. These two phenomena could also be linked: a loss of functionality of *Cas1* and *Cas2* in the MRCA of all MTC could have fostered an adaptative change in life-style of the bacterium, *i.e.* from an environmental extracellular to a host-specialized intracellular life-style. Such an hypothesis could be supported by the evolution of the CRISPR locus of *Vibrio cholerae*, with observations that the recent pandemic strains have lost their ancestral CRISPR locus (Weill, et al., 2017) and (FX Weill, personal communication). Hence, the functionality of *Cas1* and *Cas2* of MTC remains to be explored.

Regarding immunity, this study only focused on the full presence or absence of *cas* genes without exploring in detail SNP variations. As stated previously, 23/198 (12%) lacked at least part of the *cas* genes. Among these yet, all isolates still carried the *cas6, cas10/*csm1, *csm2*, and *csm3* genes. This observation matches that made previously on CRISPR clinical isolates (Freidlin, et al., 2017). Cas6 protein is involved in pre-crRNA processing. Cas10/Csm1 and Csm3 are the enzymes responsible for the catalytic activity of the crRNP (Kazlauskiene, et al., 2017; Samai, et al., 2015). Hence, regarding immunity, even if the spatial structure of the crRNP may be impaired by the absence of *csm4* and/or *csm5* in some isolates, it could remain possible that immunity occurs in all MTC isolates through the consecutive actions of Cas6 to process pre-crRNA and of Cas10/Csm1 and Csm3 to degrade DNA and/or RNA. The fact that none of the spacer is conserved in all isolates implies that, if immunity occurs, it does not always target the same DNA and/or RNA sequences.

### Global Implication of CRISPR diversity for the understanding of MTC clinical isolates evolution

In MTC, the CRISPR locus is a likely witness of a previous yet unknown evolutionary history of phage DNA invaders defense, whereas IS*6110* is a specific MTC element that belongs to the IS3 family that, through transposition, also plays a permanent role in shaping MTC genomes (Thabet and Souissi, 2017). The link between the two in evolutionary genomics remains poorly investigated until now. MTC genome actually contains a lot of other IS and transposases (88 genes retrieved in mycobrowser, (https://mycobrowser.epfl.ch/) such as IS*1081*, IS*1533*, IS*1547*, IS*1560*), but IS*6110* is the one with the largest number of copies in most isolates and especially in the reference isolate H37Rv (Cole, et al., 1998). IS*1547* was previously shown to play a role in MTC evolution however it remains poorly investigated (Fang, et al., 1999). IS*6110*-RFLP was the golden standard to define epidemiological clusters at the end of the nineties and stayed so during around 20 years, until it was replaced by MIRU-VNTR^1^ and more recently by Whole-Genome-Sequencing (Schurch, et al., 2010; Supply, et al., 2006; van Embden, et al., 1993; van Soolingen, et al., 2007) (for a recent review on evolution of TB molecular epidemiological methods, see also (Garcia De Viedma and Perez-Lago, 2018)). Previous results on IS*6110* insertion sites have shown that independent IS*6110* copy acquisition through transposition into *hot-spots* was a common mechanism explaining convergence in IS*6110* copy number in some of the MTBC sublineages (Dale, et al., 2003; Roychowdhury, et al., 2015). A recent paper on the micro- and macro-evolution of Lineage 2 of MTC in relation to IS*6110* transposition also stress the interest of such studies using WGS (Shitikov, et al., 2019). The role of the *ipl* (Insertion Preference Locus) was also stressed long time ago and showed consequences on the CRISPR locus (Fang, et al., 1999; Fang, et al., 1999; Fang and Forbes, 1997), however no generalized observations on IS-CRISPR genomics dynamics had been done so far before this study.

## Conclusions

Our study, by providing an *in-depth* reconstruction of the CRISPR locus of MTC using short reads on around 200 genomes, in combination with IS*6110*, improves our knowledge on the structure of the CRISPR locus and sheds new light on the general evolutionary mechanisms acting on MTC genomes through a first yet quantitatively limited analysis that combines CRISPR-IS combined evolutionary dynamics. By unveiling an unexpected genetic diversity of the CRISPR Locus on MTC, our study opens the way to new in-depth congruence analysis between SNP-based and repetitive sequence based MTC phylogenies. Such deeper knowledge on the natural history of tuberculosis will help us deciphering the most important key evolutionary events that shaped today’s global and local MTC genomes population structure.

## Declarations

### Ethics approval and consent to participate

N.A. This study only uses publicly available data

### Consent for publication

All authors read and accepted the final submitted version

### Competing interests

The authors declare no competing interest

### Funding

This study was funded by CNRS (Centre National de la Recherche Scientifique), The University of Paris-Saclay and the University of Bourgogne Franche-Comté through recurrent research support to the research teams.

### Authors’ contributions

CG, GR, CS conceived the study. CG developed the pipeline, GC,GR,CS chose the genomes to be analyzed, GR and CG analyzed results helped by CS; GR, CS and CG wrote the manuscript, GR drew the Figures and built the Supplementary Tables;

## Acknowledgements

Laura Morel, Valentin Pohyer, Matthieu Petrou, three previous undergraduates students who contribute to the start of the MTC CRISPR genome project are warmly acknowledged

## Data and Material availability

All genomic data used were extracted from Public genome databases (NCBI or ENA archives). Computer Program specifically developed in this paper will be made freelu available upon request to Christophe Guyeux.

## Supplementary files

Supplemental file 1 (doc) - Sequences of interest in CRISPR-Cas region of *Mycobacterium tuberculosis* complex.

Supplemental file 2 (tab) – CRISPR reconstructions highlighting 1) global structure and position of IS6110 insertions [‘IS6110’ sheet]; 2) spacer variants [‘spacer’ sheet]; 3) DR variants [‘DR’ sheet]; 4) Duplicated DVR [‘Duplic’ sheet].

Supplemental file 3 – Exploration of read numbers for the reconstruction and identification of duplications, the case of ERR718197.

Supplemental file 4 – Confirmation of sp35 presence after spacer 41 in two Sequence runs from clinical isolatess belonging to L5 and L2 respectively

Supplemental file 5 - Spacer 4, spacer 6 and spacer 38 variants in parallel with 43-spacers spoligotyping probes

Supplemental file 6 – Cumulative punctual variant numbers 5DR variants + spacer variants) in groups of 5 successive DVR from DVR1-5 to the last three DVR (DVR66-68)

## Footnotes

1 Mycobacterial Interspersed Repetitive Units-Variable Number of Tandem Repeats Typing

