## Supplementary file 1 for "Unexpected diversity of CRISPR unveils some evolutionary patterns of repeated sequences in *Mycobacterium tuberculosis*"

**Supplementery file 1 – Sequences of interest in CRISPR-Cas region of *Mycobacterium tuberculosis* complex.**

DR0: GTCGTCAGACCCAAAACCCCGAGAGGGGACGGAAAC,

DR1: GTCGTCAGACCCAAAACCCCGAGAGGGGACGGGAAC,

DR2: GTCGTCAGACCCAAAACCCCGAGAGAGGACGGAAAC,

DR3: GTCGTCAGACCTAAAACCCCGAGAGGGGACGGAAAC,

DR4: GTCCTCAGACCCAAAACCCCGAGAGGGGACGGAAAC,

DR5: GTCGTCAGACCCAAAACCACGAGAGGGGACGGAAAC,

DR6: GTCGTCGGACCCAAAACCCCGAGAGGGGACGGAAAC,

DR7: GCCGTCAGACCCAAAACCCCGAGAGGGGACGGAAAC,

DR8: GTCGTCCGACCCAAAACCCCGAGAGGGGACGGAAAC,

DR9: GTCGGCAGACCCAAAACCCCGAGAGGGGACGGAAAC,

DR10: GTCGTCAGACCCAAAACGACGAGAGGGGACGGAAAC,

DR11: GTCGTCAGGCCCAAAACCCCGAGAGGGGACGGAAAC,

DR12: GTCGTCAGACCCAAAACCCCGAGAGGGGACGGACAC,

DR13: GTCGTCAGACCCAAAACCCCAAGAGAGGACGGAAAC,

DR14: GTCGTCAGACCCAAAACCCCGAGAAGGGACGGAAAC,

DR15: GTCGTCAGACCCAAAACCCCGAGAGGGGACGTAAAC,

DR16: GTCGTCAGACCCAAAACCCCGAGAGGGGACGGAAAG,

DR17: GTCGTCAGACCCAAAACCCCGAAAGGGGACGGAAAC,

DR18: GTCGTCAGACCCAAAACCCCGAGAGGGTACGGAAAC,

DR19: GTCGTCAGACCCAAAAGCCCGAGAGGGGACGGAAAC,

DR20: GTTGTCAGACCCAAAACCCCGAGAGGGGACGGAAAC,

DR21: GTCGTCAGACCCAAAACCCCGATAGGGGACGGAAAC,

DR22: GTCGTCAGACCCAAAACTCCGAGAGGGGACGGAAAC,

DR23: GTCGTCAGACCCAAAACCCCGAGAGGGGAAGGAAAC,

DR24: GTCGTCAGACCCAAAACCCCGAGATGGGACGGAAAC,

DR25: GTCGTCAGACTCAAAACCCCGAGAGGGGACGGAAAC,

DR26: GTCGTCAGACCCTAAACCCCGAGAGGGGACGGAAAC,

DR27: GTCGTCAGACCCAAAACCCCGAGGGACGGAAAC,

DR28: GTCGTCAGACCCAAAACCCCGAGAGGGGACTGAAAC,

DR29: GTCGTCAGACCCAAACCCCGAGAGGGGACGGAAAC,

DR30: GTCGTCAGACCAAAAACCCCGAGAGGGGACGGAAAC,

DR31: GTCGTCAGACTGAGTCGTCAGACCCAAAACCCCGAGAGGGGACGGAAAC,

DR32: GTCGTCAGACCCAAAACCCCGAGACGGGACGGAAAC,

DR33: GTCTTCAGACCCAAAACCCCGAGAGGGGACGGAAAC,

DR34: GTCGTCAGACCCAAAACCCCGAGAGTGGACGGAAAC,

DR35: GTCGTCAGACCCAAAACCCCGAGAGGGGACGGAAAT,

DR36: GTCTTCAGACCCAAAACCCCGAGAGAGGACGGAAAC,

DR37: GTCGTCAGACCCAAAACCCCTAGAGGGGACGGAAAC,

rDRa1: GTCGTCAGACCCAAAACCC,

DRb1: CCCCGAGAGGGGACGGAAAC,

DRb(1): AACCGAGAGGGGACGGAAAC,

DRb2: AAAACCCCGAGAGGGGACGGAAAC,

DRb3: CCCTGAGAGGGGACGGAAAC,

esp1: TTAAAACCGTGTTGCACTGCAACCCGGAATTCTTGCAC,

esp2: CATAGAGGGTCGCCGGCTCTGGATCACGCTCCCCTAGTCGT,

esp2(1): CATAGAGGGTCGCCGGCTCCGGATCACGCTCCCCTAGTCGT,

esp3: TTTTTGCCTCATGCTTGGGCGACAGCTTTTGACCAA,

esp4: TCGCAAGCGCCGTGCTTCCAGTGATCGCCTTCTA,

esp4(1): TCGCAAGCGCCGTGCTTCCAGTGATCACCTTGTA,

esp4(2): TCGCAAGCGCCGTGCTTCCAGTGATCACCTTCTA,

esp5: TCGCGGCGCGGCATGGCACGGCAGGCGTGGCTAGGGG,

esp6: ATGTGCGCCGTCGCCGTAAGTGCCCCACGGCCCGT,

esp6(1): ATGTGCGCCGTCGCCGTAAGTACCCCACGGCCCGT,

esp6(2): ATGTGCGCCGTCGCCGTAAGTACCCCACGGCCAGT,

esp7: ATTTCGACGACAATTCGTTGACCACGAATTTTCAGA,

esp8: ACATCCCACGCGTTACCGCTGGCGCGCATCATTCATCGA,

esp9: CCATATCGGGGACGGCGACGCTGCGAGAGGACACGCCGA,

esp9(1): CCTTATCGGGGACGGCGACGCTGCGAGAGGACACGCCGA,

esp9(2): CCATATCGGGAACGGCGACGCTGCGAGAGGACACGCCGA,

esp10: TACACCACGCGTCGTGCCATCAGTCAGCGTCCTCCTC,

esp11: TTGAACACGGAGCCGTGCACATGCCGTGGCTCAGGGGT,

esp12: AACACCTCAGTAGCACGTCATACGCCGACCAATCATCAG,

esp12(1): AACACCTCAGTAGCACGTCATACGCCAACCAATCATCAG,

esp12(2): AACACATCAGTAGCACGTCATACGCCGACCAATCATCAG,

esp13: TTTTCTGACCACTTGTGCGGGATTAGCGGGCTTAG,

esp14: ACCAATGCGTCGTCATTTCCGGCTTCAATTTCAGCCT,

esp15: CTGAGGAGAGCGAGTACTCGGGGCTGCCGTCTGCGCTG,

esp15(1): CTGAGGAGAGCGAGTACTCGGGGCTGCCGTCTGCGGTG,

esp16: ACGACGTTAGGGCATGCAGCATGCCGTCCCCGTTTTTGA,

esp16(1): ACGACGTTAGGGCATGCAGCATGCCGTCCCCCGTTTTTGA,

esp17: TGCTCTTGAGCAACGCCATCATCCGGCGCCGCAGCTCCGC,

esp18: GCGTGAAACCGCCCCCAGCCTCGCCGGGGCCGCCTAG,

esp19: ACTCGGAATCCCATGTGCTGACAGCGGATTCGCAT,

esp20: CGGGCAGCGTTCGACACCCGCTCTAGTTGACTTCCGG,

esp20(1): CGGGCAGCGTTCGACACCCGCTCTAGTTGACGTCCGG,

esp21: CAGGTGAGCAACGGCGGCGGCAACCTGGCGGCCACGGGTCG,

esp21(1): CAGGTGAGCAACGGCGGCGGCAACCTGGCGGCCCCGGGTCG,

esp22: ATGGGATATCTGCTGCCCGCCCGGGGAGATGCTGTCCGAG,

esp23: TTCGTCGACCATCATTGCCATTCCCTCTCCCCACGT,

esp24: TTGCGCCAACCCTTTCGGTGTGATGCGGATGGTCGGCTCGG,

esp24(1):TTGCGCCACCCTTTCGGTGTGATGCGGATGGTCGGCTCGG,

esp25: CTTGAATAACGCGCAGTGAATTTCGCGGATCAGACCC,

esp26: ATTCGCACGAGTTCCCGTCAGCGTCGTAAATCGCCA,

esp26(1): ATTCGCACGAGTTCCCTTCAGCGTCGTAAATCGCCA,

esp26(2): ATTCGCACGAGTTCCCGTCAGCGTCTTAAATCGCCA,

esp27: CCGGCAACAATCGCGCCGGCCCGCGCGGATGACTCCG,

esp28: CGCATGGACCCGGGCGAGCTGCAGATGGTCCGGGAG,

esp29: TGGATTGCGCTAACTGGCTTGGCGCTGATCCTGGTG,

esp29(1): TGGATTGGGCTAACTGGCTTGGCGCTGATCCTGGTG,

esp30: TCCACATCGATTTCCTTGACCTCGCCAGGAGAGAAGATCAC,

esp31: TCGTCGACGATCGCGTCGATGTCGATGTCCCAATCGTCGA,

esp31(1): TCGTCGACGATCGTGTCGATGTCGATGTCCCAATCGTCGA,

esp32: TTGGAGCGTGTCACCGCAGACGGCACGATTGAGACAA,

esp33: CCTCAGCTCAGCATCGCTGATGCGGTCCAGCTCGTCCGT,

esp33(1): CCTCAGCTCCGCATCGCTGATGCGGTCCAGCTCGTCCGT,

esp33(2): CCTCAGCTCAGCATCGTTGATGCGGTCCAGCTCGTCCGT,

esp34: CCAACCTCACCGCCTGCTGGGTGAGACGTGCTCGCCGCGA,

esp35: TCGGGGAGCCGATCAGCGACCACCGCACCCTGTCA,

esp35(1): TCGGGGAGCCGATCAGCGACCACCGCACCGTCA,

esp36: CTTCAGCACCACCATCATCCGGCGCCTCAGCTCAGCAT,

esp37: CCTTCGACGCCGGATTCGTGATCTCTTCCCGCGGATAG,

esp38: TGCCCCGGCGTTTAGCGATCACAACACCAACTAATG,

esp38(1): TGCCCCAGCGTTTAGCGATCACAACACCAACTAATG,

esp39: CAGCGAAATACAGGCTCCACGACACGACCACAACGC,

esp40: TCTTGACGATGCGGTTGCCCCGCGCCCTTTTCCAGCC,

esp41: AGGTTCGCGTCAGACAGGTTCGCGTCGATCAAGTCCG,

esp42: TTTATCACTCCCGACCAAATAGGTATCGGCGTGTTCAA,

esp42(1): TTTATCACTCCCGACCAAATAGATATCGGCGTGTTCAA,

esp43: TCGACACCGACATGACGGCGGTGCCGCACTTGACGCA,

esp44: CTTTGCGAAGTCACCTCGCCCACACCGTCGAAGCGCCT,

esp45: GCGGATGGTGGGCAAGTTGGCGCTGGGGTCTGAGTCAA,

esp46: CTGCATCCGGAAAGTCCGTACGCTCGAAACGCTTCCAACGT,

esp47: TCGAAATCCAGCACCACATCCGCAGCTGCGGCATGCTCCCGAA,

esp48: GCGAGGAACCGTCCCACCTGGGCCTGCCCCCAGCGG,

esp49: TCAATAACACTTTTTTTGAGCGTGGCGCGGTTGAGAGT,

esp50: ACGGAAACGCAGCACCAGCCTGACAATCTTATTCTCGC,

esp50(1): ACGGAAACGCAGCACCAGCCTGACAATCTTATTCACGC,

esp51: ATTTTGAGCGCGAACTCGTCCACAGTCCCCCTTTCAG,

esp51(1): ATTTTGAGCGCGAACTCGTCGACAGTCCCCCTTTCAG,

esp52: GCCCCGTGGATGGCGGATGCGTTGTGCGCGCAAGT,

esp53: CCGACGATGGCCAGTAAATCGGCGTGGGTAACCGATCCGG,

esp54: CCTCGCAGAAAAGGGCATCGATCATGAGAGTTGCGTTGAT,

esp55: AATGCTGGCGACGATTTTCGCTGTTGTGGTTCTCATT,

esp56: GCACACCAGCACCTCCCTTGACAATCCGGCAGATCAGAC,

esp56(1): GCACACCAGCACCTCCCTTGGCAATCCGGCAGATCAGAC,

esp57: TCGCGGGCTCTGGCCTAAGGGTGCTGACTTCGCCTGTA,

esp57(1): TCGCGGGCTCTGGCATAAGGGTGCTGACTTCGCCTGTA,

esp57(2): TCGCGGGCTCTGGTCTAAGGGTGCTGACTTCGCCTGTA,

esp58: CCGACGACGAGCAGCGGCATACAGAGCCACGGATACGCCAG,

esp59: TTGCATCCACTCGTCGCCGACACGGCGGACTTCCGCGA,

esp60: TGGTAATTGCGTCACGGCGCGCCTGGCGGGCCGATT,

esp60(1): TGGTAATTGCATCACGGCGCGCCTGGCGGGCCGATT,

esp60(2): TGGTAATTGCGTCACGGCGCGCCTGGCGGGCCTATT,

esp61: ACCATCCGACGCAGGCACCGAAGTCGATGACAAGCC,

esp62: TAGTACGCCATCTGTGCCTCATACAGGTCCAGTGCCCT,

esp63: CTGACGGCACGGAGCTTTCCGGCTTCTATCAGGTA,

esp63(1): CTGACGTCACGGAGCTTTCCGGCTTCTATCAGGTA,

esp63(2): CTGACGGCACGGAGCTTTCCGGCTTCTATTAGGTA,

esp64: CCTCATGGTGGGACATGGACGAGCGCGACTATCGGG,

esp64(1): CCTCATGGTGGGACATGGACGAGCTCGACTATCGGG,

esp65: TGGACGCAGAATCGCACCGGGTGCGGGAGGTGCAGCA,

esp66: GCATATCGCCCGCCACACCACAGCCACGCTACTGCTCCAT,

esp66(1): GCATATCGCCCACCACACCACAGCCACGCTACTGCTCCAT,

esp67: ACACCGCCGATGACAGCTATGTCCGAGTGACATCCTCCCA,

esp68: TTGAACCGCCCTTCGCGCGGTGTTTCGGCCGTGCCCGA,

esp69: TTGGAGGTGTCGTTGCATTCGTCGACTGCGTGGTATT,

esp70: ATGCCCGCTGGTAGCGGCCCCGGCGGCGCCGAGAA,

esp71: TTGGTGATCTTCCATGACTTGACGCCGTTGACCGCGAT,

esp72: GCGGTGCTCGATGCGGCCACTAGGCCGCGGCCAACTCGG,

esp73: TCGGTGCTGACCCCATGGATGCGAAGCGTAGACCTCTGT,

esp74: CAACAAGGTCTACGCGTCGAGGTCCACGGCTCAGAA,

esp75: ATGTGGCGATTACGCCTGATCAGGCGAAGGCGA,

esp76: TTCAGTAAATTGCAGCGACGGGCGAATCCAGCAACC,

esp77: CTTCAACGACGCTGTATTGGGCCATGTGGGCTGGTCTTTCA,

esp78: AGCAGCATGGACGGTTTCGCCTGTATCCGGGTAAAAAA,

esp79: TCGATGGCGGCGCGGTTGCGGATGTGGTGGTCGCGTAGC,

esp80: TTGGCGTACATAGCGAGCTGTGCGGCAGTAGAGTGCGG,

esp81: CTTCGATTGTGCCGCCGCGGGTTTCGTTCAC,

esp82: TCCCCGGGCTGGGGCGTGTGTTCGTAGTCGCCTAAT,

esp82(1): TCCCCGGGCTGAGGCGTGTGTTCGTAGTCGCCTAAT,

esp83: CCTTGCCAAGGGAGGTGCTGGTGTGCTTATGCCTAACAG,

esp84: TAGTTGACGCCAAATGTTTGGACTGTGATCAATTCAA,

esp85: GCTTGACAATGCGGTTGTCGCGCGCCTTTTTCCAGCCGAG,

esp86: GTGTGTTTCAACGGGTTTCAGTTTTCTTGTCCCAGTG,

esp87: ACTGGTTGTTGCCCGGCGACGGCGGCGCGTTGGCGTGTC,

esp88: CATCCAGAGGTCGAAGTGGTGTTCGGTGTTCTCCTGTAC,

esp89: TTCTGCGCGGCGATCCGGTCATGACGAGCCCGCAG,

esp90: ATCACGACACGGCCTGATCGGTGTCGGTGGCGGCAAGTCGTCAGA,

esp91: CAGAAGGGTAGCGTCGGCCGGATTGTCTGGCCCAC,

esp92: CATCGAACAAGCGCGCGTCGGCTAAGCACGCGTCTGTCAA,

esp93: TCTCCGTGATAGGTGAGGACCACCGAATCACCATCA,

esp94: TCTGGTAGTGGGCTTCTGCCGGTGCGCGGGTTTGTGTCT,

esp95: CCGCGTCATCGGGCACGACCACGAACAGGTCCGTGAAG,

esp96: ATACCGGAAGCACATTCGCGCCAGTGGTCGTCGCAGAAG,

esp97: ATTCCGACTAGCGCGTCGTCGTTTCCAGCCTCAAT,

esp98: CGACAAGTTTGCGCGGATCAAGTCCGCGCCGGTCAA,

IS6110:TGAACCGCCCCGGCATGTCCGGAGACTCCAGTTCTTGGAAAGGATGG,

IS6110c:TGAACCGCCCCGGTGAGTCCGGAGACTCTCTGATCTGAGACCTCAGC,

IS6110c(1):TGAACCACCCCGGTGAGTCCGGAGACTCTCTGATCTGAGACCTCAGC,

finIS6110c:AAGAACTGGAGTCTCCGGACATGCCGGGGCGGTTCA,

finIS6110:AGATCAGAGAGTCTCCGGACTCACCGGGGCGGTTCA,

finIS6110*DRb1: GACTCACCGGGGCGGTTCACCCCGAGAGGGGACGGAAAC,

rDRa1*IS6110:GTCGTCAGACCCAAAACCCTGAACCGCCCCGGCA,

Csm4[:341]*IS6110c: GATGGCACGGCCGACCTGAATGAACCGCCCCGGTGAGTC,

motif_debut1: CATCATCAGCAGGCATTGTTACCACACGCTGGACGAATTGTCCATAGA,

motif_debut2: CATCATCAGCAGGCATTGTTACCACACGCTGGACGAATTGTCCATCGA,

motif_debut3: CATCATCAGCAGGCATTGCTACCACACGCTGGACGAATTGTCCATAGA,

motif_debut4: CATCATCAGCAGGCATTGTTACCACACGCCGGACGAATTGTCCATAGA,

motif_debut5: CATCATCAGCAGGCATTGTTACCACACGCTGGCCGAATTGTCCATAGA,

motif_fin1:TACGACGACTGGGTCGCCACCGCGTCTGTTGACCGGCATTCAGGATGA,

motif_fin2:GCATGATGGCGGCGTTGACGGTGAGGACGTTTGGTCATGAAATGA,

motif_fin3:CCCCGCCGGGAGATGTCCGGCGGGGTCGGTGGTGTTCGGGGTGTCGGTGTGGTGT,

Rv2816c: GCTTGTCAGCGCAGAGGAGTTTGTGTTCTTTTGA,

Rv2813c: TCAGTCTGCCGTGACTTCGGCGATGGCGG

Cas6_debut: TTGGCTGCTCGCCGAGGCGGCATCAGAAGGACGGATCTGCTACGTCGGAGTGGCCAGCC,
Cas6_debut2: TTGGCTGCTCGCCGAGGCGGCATCAGAAGGGCGGATCTGCTACGTCGGAGTGGCCAGCC,
Cas6_debut3: TTGGCTGCTCGCCGAGGCGGCATCAGAAGGACGGATCTGCTACGTTGGAGTGGCCAGCC,
Cas6_fin: TGGGCGCGATCCGGGTCCAGCCACTGGCACCGAGGGAAAAATGCGTACCGAAGCCATGA,
Csm1_debut: ATGAACCCGCAACTCATCGAGGCCATAATCGGCTGCCTCTTGCACGACATTGGCAAACC,
Csm1[:2312]*Csm3[247:]:CTCACGCGCATGCGTAACCCCACCGGTGACACAGCGCCCATATCCGTCGGCTTTTCGGC,
Csm1_fin: AACTCAAGACCGCGCTGCACCTCTACATCTATCGCACTCGCAAGGAGGAGTCCGAATGA,
Csm1_fin2: AACTCAAGACCGCGCTGCACCTCTACATCTATCGCACTCGCAAAGAGGAGTCCGAATGA,
Csm2_debut: ATGAGCGTCATCCAAGACGACTATGTGAAACAGGCCGAAGTAATTCGCGGCCTGCCAAA,
Csm2_debut2: ATGAGCGTCATCCAAGACGACTATGTAAAACAGGCCGAAGTAATTCGCGGCCTGCCAAA,
Csm2_fin: GCCGGTACATGGAAGCCCTAGCCGCATACAAGAAGTACCTCGATCCGAAGGACAAGTGA,
Csm3_debut: ATGACTACGAGCTACGCCAAGATCGAGATAACCGGGACACTGACCGTCCTGACGGGCCT,
Csm3[:433]*IS6110c: GGGTGACCGCAAAGGCAAACCTTCGCCAGATGGAACGCGTGATCTGAACCGCCCCGGTG,
Csm3_fin: TCGGCGCCCTCGACGGTTCTCTGCTGGAGAAGCTAAACCATGAACTCGCGGCTGTTTAG,
Csm4_debut: ATGAACTCGCGGCTGTTTAGGTTCGACTTCGACCGCACACACTTCGGCGACCACGGCCT,
Csm4[:245]*IS6110: CCCGATTACCTGGTTCCCAAGCCCCTGCACAGCGTTCGGTCTGAACCGCCCCGGCATGT,
Csm4[:341]*IS6110c: CAGCTTGGCAGCTTCCTCGATGGCACGGCCGACCTGAATGAACCGCCCCGGTGAGTCCG,
Csm4_fin: ATCCGGTCTACAGCTACGCGCGACCGCTATTTCTCGCACTCCCGGAGTCCGCCGCATGA,
Csm5_debut: ATGAACACCTACCTGAAGCCGTTCGAACTCACGCTGCGGTGCCTGGGGCCGGTGTTTAT,
Csm5[:527]*IS6110: TCCGGGACACCAGACGCGGGAGCACCGGCAGTACGGTGAACCGCCCCGGCATGTCCGGA, #TGAACCGCCCCGGCATGTCCGGAGACT
Csm5[:647]*IS6110: GCGATCAGGGTCACCGACTCACCTGCACTGAGAACAAGTGAACCGCCCCGGCATGTCCG,
Csm5[:837]*IS6110: GGGCGAGCGGTTCCTTGAAACGCTGGCCGAGACAGCCGCGTGAACCGCCCCGGCATGTC,
Csm5[:839]*IS6110c: GGCGAGCGGTTCCTTGAAACGCTGGCCGAGACAGCCGCGTCTGAACCGCCCCGGTGAGT, #GGGCGAGCGGTTCCTTGAAACGCTGGCCGAGACAGCCGCGTTGAACCGCCCCGGTGAGTCCGGAGAC
Csm5[:839]*IS6110: GGCGAGCGGTTCCTTGAAACGCTGGCCGAGACAGCCGCGTCTGAACCGCCCCGGCATGT,
Csm5_fin: TCGACAACATATGCTACGAGATGGGTCAGTGCGAGCTGTCGATCAGGAGAGCCGAATGA,
Csm6_debut: GTGCTATTCCTCAGCGCCGAGATAGCTGCCTTTGAGAACGCGGACCGGCGGTACTCCGC,
IS6110*Csm6[294:]: GTCTCCGGACTCACCGGGGCGGTTCAAAGCACGCCTGCCCGGGCATTGAGCAAGCCTGG,
IS6110c*Csm6[293:]: TCTCCGGACATGCCGGGGCGGTTCATAAGCACGCCTGCCCGGGCATTGAGCAAGCCT,
Csm6[:439]*IS6110c: CCCCAACCGTTGCTTTGAGGCGACTTCCGCTGCGCTCGGCGTGAACCGCCCCGGTGAGT,
Csm6[:938]*IS6110: CTGCTCCGCCAATTCGCACCCGATCGAGTTGGTGCTCTTGAACCGCCCCGGCATGTCCG,
Csm6[:1021]*IS6110: CACGAGATCGTCTCAATCAGTGAGGATCGCATCACGATGAACCGCCCCGGCATGTCCGG,
Csm6_fin: TCTATGACCGGTTGAACGACGAGATCATCCGGCAGATTGATATGGCACCGCTGGGCTAA,
Cas1_debut: ATGGTGCAGCTGTATGTCTCGGACTCCGTGTCGCGGATCAGCTTTGCCGACGGCCGGGT,
Cas1_debut2: ATGGTGCAGCTGTATGTCTCGGACTCCGTGTCGTGGATCAGCTTTGCCGACGGCCGGGT,
Cas1[:800]*esp19[16:]: GTCGACACCCGGGCTTTCAGCAAGAACTCCGACACGGGTGCTGACAGCGGATTCGCAT,
Cas1_fin: CCGGGCACCCGTCGCGGCTCGTCGATATCGATATCACCTCCGAGCCATCCGGAGCCTAA,
Cas1_fin2: CCGGGCACCCGGCGCGGCTCGTCGATATCGATATCACCTCCGAGCCATCCGGAGCCTAA,
Cas2_debut: ATGCCCACTCGCAGCCGTGAGGAGTACTTCAATCTCCCGCTCAAAGTGGACGAGTCCAG,
Cas2_fin: CAGTTACGTTCTACGGAAGGGGACGGCTTGTCAGCGCAGAGGAGTTTGTGTTCTTTTGA,
motif_debut: CATCATCAGCAGGCATTGTTACCACACGCTGGACGAATTGTCCATAGA,
motif_debut*DR0: GACGAATTGTCCATAGAGTCGTCAGACCCAAAACCCCGAGAGGGGACGGAAAC,
DR0*esp1: GAGAGGGGACGGAAACTTAAAACCGTGTTGCACTGC,
motiffin(1)_debut: TACGACGACTGGGTCGCCACCGCGTCTGTTGACCGGCATTCAGGATGAGCATGATGGCG,
motiffin(2)_debut: TACGACGACTGGGTCGCCACCGTGTCTGTTGACCGGCATTCAGGATGAGCATGATGGCG,
motiffin_milieu: GCGTTGACGGTGAGGACGTTTGGTCATGAAATGACCCCGCCGGGAGATGTCCGGCGGGG,
motiffin_fin: TCGGTGGTGTTCGGGGTGTCGGTGTGGTGT,
Rv2812c_debut: TCATTTCGGTTGGGCCTCCTGGCCGGTGAGCAGGTCGAGCTGGCCGGGGATCTGATCGG,
Rv2812c_fin: AACCCGATCGCCCTCGCGCGCTCCGCGCGCACCTTCTCCTCGTCATCGCCGACCGCCAC,
Rv2811c_debut: CTACGGCAGGGTCGACTCGTGTTGCACCCACTCGCCGGGCCAGCCCGGCGCCAACAACC,
Rv2811c_fin: AGCTCACCGGCCGCCAGCCGACGCTCGACTTGATCGACATCTGCTTCTACGGTGACCAC,
Rv2809c_debut: TCATCGGTTCCACTCCACCAGCTTCATCCGCACGTCGACACCTGGTCCGAGCTTCTGCT,
Rv2809c_fin: CACCGCATCTGTGCGAGCAGTTTGGGGAGCGTCGTATCGTCCCTGGCTGCGTACGTCAT,
Rv2808c_debut: TTATCGGTATATCGGCCACGTCTCTAAGTATGCGTCAATTGCTCGTTCGATTTTCAACC,
IS6110c*Rv2808c[194:]: TCCGGACATGCCGGGGCGGTTCATTCTCTAATTCTTGCTCGATCACGGGACGGTGCTCC,
Rv2808c_fin: AATTCTTGCTCGATCACGGGACGGTGCTCCGTTGAAATAGCATCGAGAACGTTGCTCAC,
Rv2807c_debut: TTACTTCGCCTTGGCCAATCGGTTGATTGACGGTTGCAATGATTGCAGGTCGATGTGGC,
Rv2807c_fin: CCCGGGCCGGTCAGCATCCGCCGGGCCGTCGAACGACCCATCCCGGTGGTGGACACCAC
