## Supplementary file 3 for "Unexpected diversity of CRISPR unveils some evolutionary patterns of repeated sequences in *Mycobacterium tuberculosis*"

**Supplementary file 3 – Exploration of read numbers for the reconstruction and identification of duplications, the case of ERR718197 (L1.1.1.7).** This table shows the number of reads containing at least the 12 last nucleotides of one spacer at its beginning, followed by a Direct Repeat, followed by at least 12 first nucleotides of one spacer.

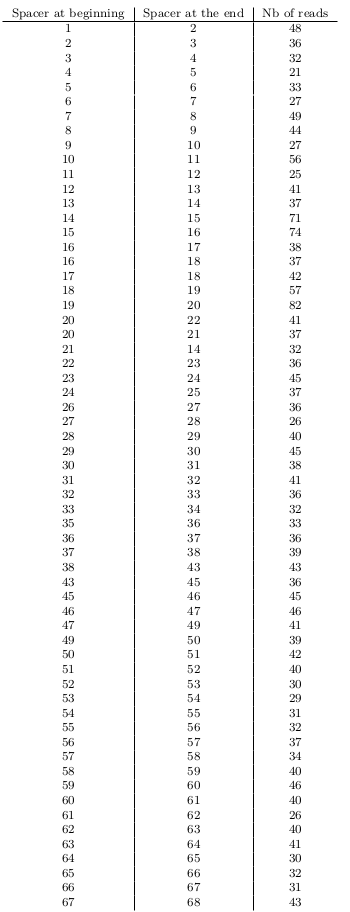
