## Supplementary file 4 for "Unexpected diversity of CRISPR unveils some evolutionary patterns of repeated sequences in *Mycobacterium tuberculosis*"

**Supplementary File 4 – Confirmation of sp35 presence after spacer 41 in two Sequence runs from strains belonging to L5 and L2 respectively.** A. Reads from SRR998631 (*M. tuberculosis* variant *africanum*) having at least 12 nucl of spacer 41, one DR0 followed by 12 nucl of sp. 35. B. Reads from ERR234248 (L2.1) having at least 12 nucl of spacer 41 and 12 nucl of sp. 35.

A

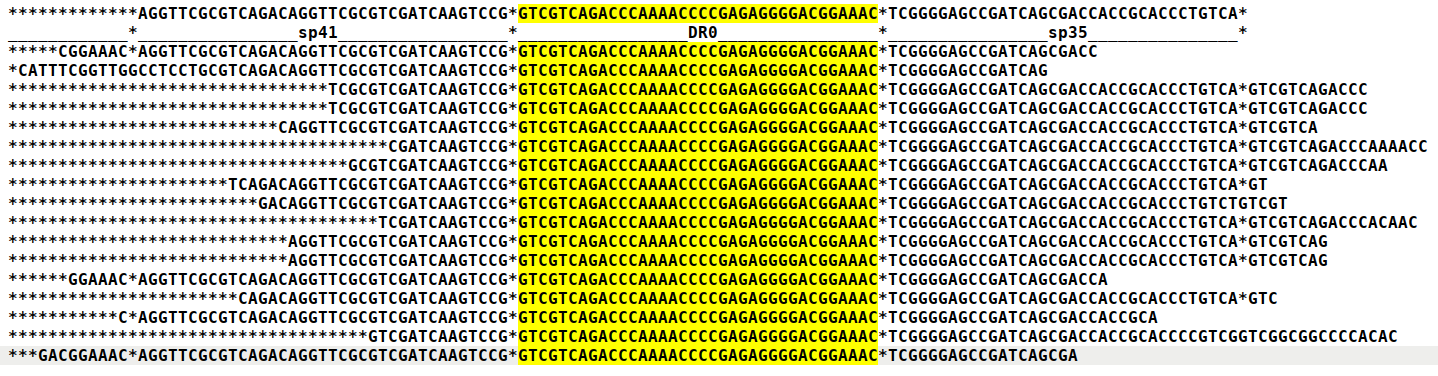

B

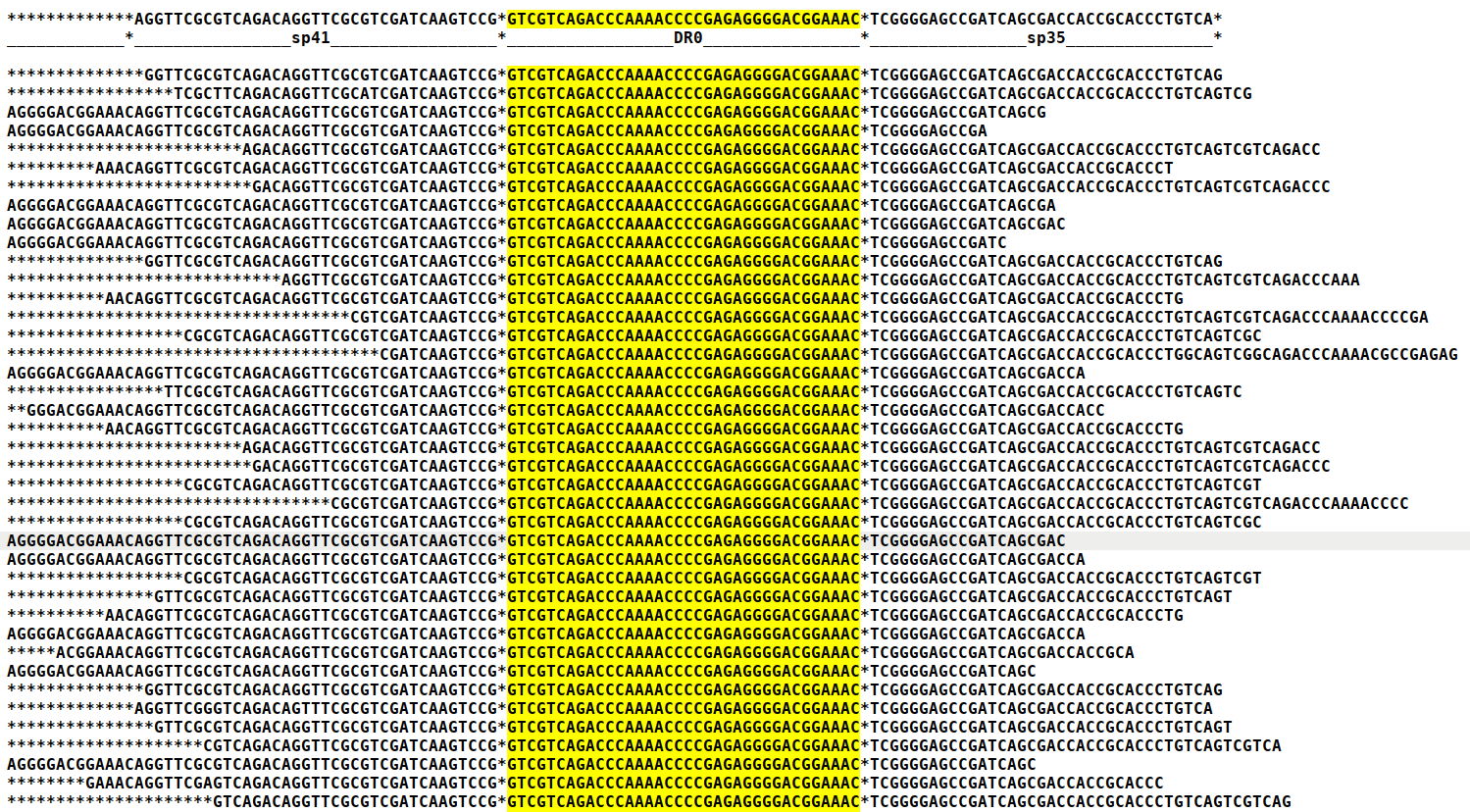
