## Supplementary file 5 for "Unexpected diversity of CRISPR unveils some evolutionary patterns of repeated sequences in *Mycobacterium tuberculosis*"

**Supplementary file 5 - Spacer 4, spacer 6 and spacer 38 variants in parallel with 43-spacers spoligotyping probes.**

A- **Spacer 4**

>most frequent sequence (esp4)

TCGCAAGCGCCGTGCTTCCAGTGATCGCCTTCTA

>variant 1 (esp4(2))	[found in all L6 strains]

TCGCAAGCGCCGTGCTTCCAGTGATC**A**CCTTCTA

>variant 2 (esp4(1))	[found in some L6 strains]

TCGCAAGCGCCGTGCTTCCAGTGATC**A**CCTT**G**TA

Spoligo43 probe #3

3…………... CCGTGCTTCCAGTGATCGCCTTCTA

B- **Spacer 6**

> most frequent sequence (esp6)

ATGTGCGCCGTCGCCGTAAGTGCCCCACGGCCCGT

> variant 1 (esp6(1))

ATGTGCGCCGTCGCCGTAAGT**A**CCCCACGGCCCGT

> variant 2 (esp6(2))

ATGTGCGCCGTCGCCGTAAGT**A**CCCCACGGCC**A**GT

C- **Spacer 38**

>esp38

TGCCCCGGCGTTTAGCGATCACAACACCAACTAATG

>esp38(1)	[found in L1.1.1 strains]

TGCCCC**A**GCGTTTAGCGATCACAACACCAACTAATG

Spoligo43 probe #28

28 .TGCCCCGGCGTTTAGCGATCACAAC
