## Supplementary figures and images for "Unexpected diversity of CRISPR unveils some evolutionary patterns of repeated sequences in *Mycobacterium tuberculosis*"

### Supplementary file 6

#### Slide 1
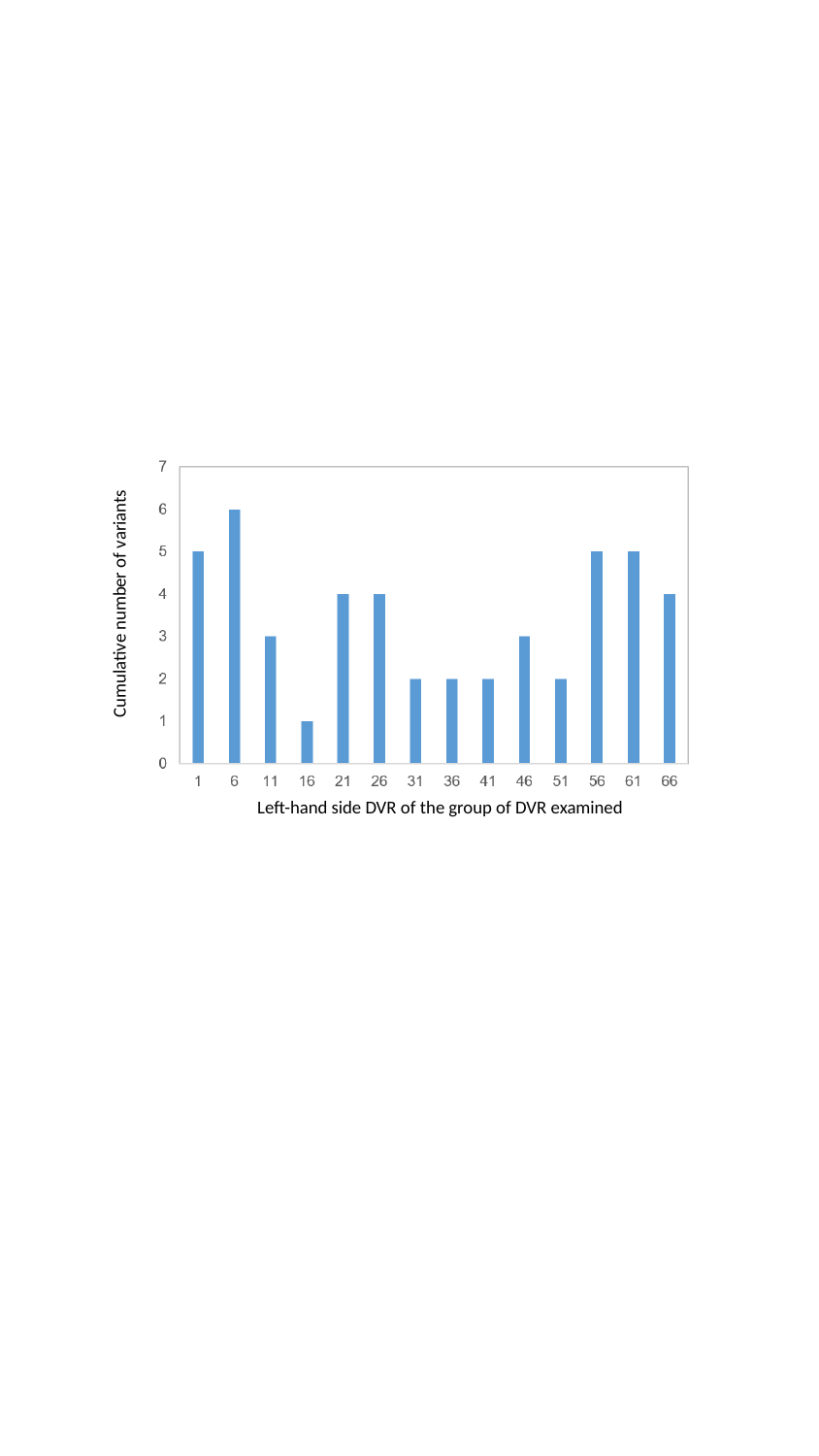

Cumulative number of variants
Left-hand side DVR of the group of DVR examined
